# Arabidopsis REM transcription factors and GDE1 shape the DNA methylation landscape through the recruitment of RNA Polymerase IV transcription complexes

**DOI:** 10.1101/2025.02.21.639493

**Authors:** Zhongshou Wu, Yan Xue, Shuya Wang, Yuan-Hsin Shih, Zhenhui Zhong, Suhua Feng, Jonathan Draper, Allen Lu, Jihui Sha, Lu Li, James Wohlschlegel, Keqiang Wu, Steven E. Jacobsen

**Author notes:** **Materials & Correspondence:** Steven E. Jacobsen.

## Abstract

DNA methylation plays crucial roles in gene regulation and transposon silencing. In plants, the maintenance of DNA methylation is controlled by several self-reinforcing loops involving histone methylation and non-coding RNAs. However, how methylation is initially patterned at specific loci is unknown. Here, we describe four Arabidopsis REM transcription factors, VDD, VAL, REM12 and REM13, that recognize specific sequence regions, and together with the protein <u>G</u>ENETICS <u>D</u>ETERMINES <u>E</u>PIGENETICS1 (GDE1), recruit RNA polymerase IV transcription complexes to generate 24-nucleotide small interfering RNAs (24nt-siRNAs) that guide DNA methylation to specific loci. In the absence of *GDE1*, Pol IV transcription complexes redistribute to sites bound by a different factor called REM8. These results suggest that REM proteins act as sequence specific DNA binding proteins that pattern siRNAs and methylation at specific sites in the genome, highlighting the role of genetic information in determining epigenetic patterns.

---

Cytosine DNA methylation is critical to silence transposable elements^1^ and maintain genome stability in eukaryotes^2,3^. DNA methylation in plants is found in three different sequence contexts: CG, CHG, or CHH (where H is A, T, C). The plant-specific RNA-directed DNA methylation (RdDM) pathway utilizes small interfering RNAs (siRNAs) to establish DNA methylation in all three contexts^4,5^. The initiation of RdDM involves the transcription of siRNA precursor RNA transcripts (25-40 nt) by RNA polymerase IV (Pol IV)^6-8^. Four related chromatin remodeling factors, CLASSY (CLSY) 1-4, are required for the recruitment of Pol IV and act as tissue-specific regulators of 24nt-siRNA expression^9,10^. RdDM plays a key role in shaping the diverse DNA methylation landscapes across various somatic tissues and cell types^9-20^. For example, in ovule tissue, siRNAs are prominently produced at siren loci, and corresponding siren DNA sites are extensively methylated. Similar patterns are seen in both monocot and dicot plants^10,15,16^. In addition, HyperTE loci, a class of highly expressed siRNA loci in male meiocyte cells, were also identified recently^12^. Although siRNAs are required for DNA methylation targeting, the mechanisms by which Pol IV is targeted to specific loci to initiate siRNA production are largely unknown.

## GDE1 is a previously uncharacterized protein localizing to RdDM sites

A previously unknown protein, AT1G77270 (hereafter named GDE1), was identified in a crosslinked immunoprecipitation-mass spectrometry (IP-MS) dataset in which chromatin was immunoprecipitated with MORC7^21,22^, a protein involved in RdDM. To determine the genomic localization of GDE1, we performed chromatin immunoprecipitation (ChIP)-seq with flower tissues from *pGDE1::GDE1-3FLAG* (*GDE1-3FLAG*). GDE1 largely co-localized with MORC7 across the genome, as well as with key components of the RdDM pathway, including Pol IV and Pol V (Extended Data Fig.1a-d). These results suggest that GDE1 may be related to RdDM function.

Motif enrichment analysis of GDE1 ChIP peaks with a fold enrichment greater than five revealed the presence of a highly conserved motif (hereafter named *clsy3 clsy4* motif1), which the RdDM component CLSY3 was previously shown to favor as well (Fig. 1a)^10^. We found that CLSY4 ChIP signals exhibited strong enrichment at CLSY3 binding sites, with over 92% of CLSY3 peaks overlapping with CLSY4 peaks (Extended Data Fig.1e-f). Furthermore, GDE1 was highly enriched at both CLSY3 and CLSY4 binding sites (Extended Data Fig.1g-h). Similar to the unique expression pattern of CLSY3 and CLSY4^10^, GDE1 was highly expressed in flower tissues (Extended Data Fig.1i) based on the ePlant database^23^. Taken together, these results suggest that GDE1 might collaborate with CLSY3/4.

**Fig. 1.**
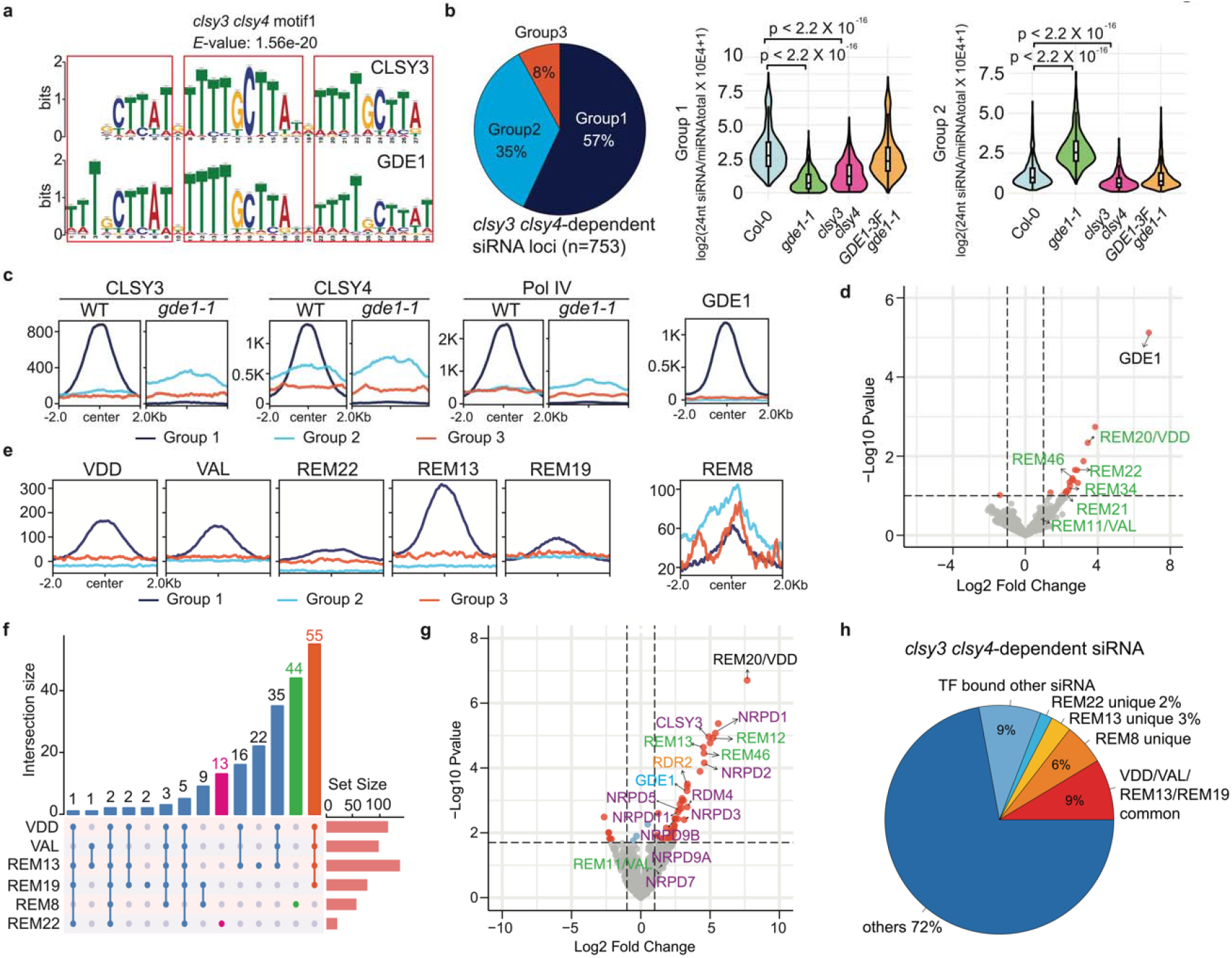
REM transcription factors associate with GDE1 and Pol IV complex at *clsy3 clsy4*-dependent siRNA loci. **a**, TOMTOM analysis showing the similarities of CLSY3 and GDE1 binding motifs. Three repeats are boxed in red. **b**, Pie chart showing the percentage of three groups of *clsy3 clsy4*-dependent siRNA sites and violin plots showing 24nt-siRNA levels in Group1 and Group2 of *clsy3 clsy4*-dependent sites. **c**, Metaplot showing enrichment of CLSY3/4 and Pol IV ChIP-seq signal over three groups of *clsy3 clsy4*-dependent sites in WT and *gde1-1* backgrounds and enrichment of GDE1 over three groups of *clsy3 clsy4*-dependent sites. **d**, Volcano plot showing proteins that have significant interactions with GDE1 as detected by IP-MS, with REM type transcription factors labeled with green letters. **e**, Metaplot showing enrichment of VDD, VAL, REM22, REM13, REM19, and REM8 ChIP-seq signal over three groups of *clsy3 clsy4*-dependent sites. **f**, UpSet plot displaying the comparative analysis among all tested transcription factors bound to *clsy3 clsy4*-dependent siRNA loci. VDD, VAL, REM13 and REM19 common targets are labeled with orange, REM8 unique target is labeled with green, REM22 unique target is labeled with magenta. **g**, Volcano plot showing proteins that have significant interactions with VDD as detected by IP-MS, with REM type transcription factors labeled with green letters, GDE1 labeled with cyane letters, CLSY3, RDM4, and Pol IV subunits labeled with purple letters. **h**, Pie chart displaying the percentage of transcription factors binding to *clsy3 clsy4*-dependent siRNA loci.

## GDE1 colocalizes with the Pol IV recruiters CLSY3 and CLSY4 at some *clsy3 clsy4*-dependent siRNA loci and is required for siRNA production

CLSY3/4 have been reported to be required for Pol IV activity and siRNA generation in ovules^9,10^. Our analysis identified 753 siRNA sites that were reduced in the *clsy3 clsy4* double mutant in ovule tissues (Fig. 1b). Among the *clsy3 clsy4*-dependent siRNA loci, 57% were reduced in *gde1-1* ovules (defined as Group 1 sites), while 35% were increased (Group 2) (Fig. 1b and Extended Data Fig.2a-d). The endogenous promoter-driven 3FLAG-tagged version of GDE1 was shown to complement the *gde1-1* phenotype in siRNA biogenesis (Fig. 1b and Extended Data Fig.2a-c). CLSY3/4 and Pol IV displayed strong enrichment at Group 1 in the Col-0 (WT), but this enrichment was largely lost in the *gde1-1* background (Fig. 1c and Extended Data Fig.2d). In contrast, CLSY3/4 and Pol IV were barely enriched at Group 2 in WT, but more ChIP signals were observed in the *gde1-1* background (Fig. 1c and Extended Data Fig.2d). Consistent with GDE1 playing a role in recruiting siRNA biogenesis to Group 1 sites, we found that GDE1 was strongly localized to Group 1 sites (Fig. 1c). These findings suggest that GDE1 plays a pivotal role in recruiting the CLSY3/4-Pol IV complex to generate siRNAs at specific *clsy3 clsy4*-dependent siRNA loci, and that CLSY3/4-Pol IV is redistributed to other loci in the absence of GDE1.

## GDE1 interacts and colocalizes with REM transcription factors

Unexpectedly, none of the CLSY3/4-Pol IV components were identified in GDE1-3FLAG IP-MS experiments (Fig. 1d and Supplementary Table 1). However, co-immunoprecipitation (co-IP) assays using F2 transgenic plants expressing GDE1-3FLAG with CLSY3-9myc or Pol IV-9myc successfully detected their association (Extended Data Fig.2e), suggesting that these interactions may be weak or transient and do not survive the IP-MS protocol.

Interestingly, multiple members from the REM (Reproductive Meristem) transcription factor family were pulled down by GDE1 (Fig. 1d, Supplementary Table 1 and Extended Data Fig.2e). Two of these REM proteins, VDD and VAL, showed strong ChIP signals over the same set of *clsy3 clsy4*-dependent siRNA loci in Group 1 (Fig. 1e-f and Extended Data Fig.3a-d). Another factor, REM22, was highly enriched at a distinct subset of *clsy3 clsy4*-dependent siRNA loci within Group 1, but also showed some colocalization with VDD and VAL (Fig. 1e-f and Extended Data Fig.3a-e).

To further explore transcription factor-GDE1 complexes, IP-MS was performed by using *VDD-3FLAG* transgenic lines. VDD successfully pulled down GDE1, CLSY3, Pol IV subunits^24^, RDR2, and RDM4, an IWR-type transcription factor known to interact with Pol IV^1^ (Fig. 1g), indicating that VDD forms a complex with GDE1 and Pol IV complex components. In addition, we detected VAL, REM46 and additional members of the REM transcription factor family (Fig. 1g), indicating that these transcription factors interact with each other *in vivo*. Consistent with this IP-MS data, VAL was previously reported to interact with itself and VDD by yeast two-hybrid (Y2H) and a bimolecular fluorescence complementation (BiFC) assay^25^. Furthermore, previous DAP-seq data from REM12, one of the transcription factor proteins identified from the VDD-3FLAG IP-MS, showed a similar motif enrichment as the *clsy3 clsy4* motif1^26^ (Extended Data Fig.3f), and REM12 DAP-seq locations highly overlapped with the same set of *clsy3 clsy4*-dependent siRNA loci bound by VDD and VAL (Extended Data Fig.3g). Additionally, most of these REM transcription factors showed similar expression patterns as CLSY3/4 and GDE1, exhibiting high expression in ovules in comparison to other tissues (Extended Data Fig.3h). Taken together, these results suggest that multiple REM transcription factors may be involved in the recognition of the *clsy3 clsy4*-dependent siRNA loci and their association with GDE1 likely recruits CLSY3/4-Pol IV to generate siRNAs for locus-specific DNA methylation.

To search for additional REM transcription factors that might recognize the *clsy3 clsy4*-dependent siRNA loci, we performed ChIP-seq using 9myc-tagged REM transcription factors identified in the IP-MS data (Fig. 1d, 1g, Supplementary Table 1 and 2) and/or co-expressed with GDE1 by ATTED-II RNA coexpression analysis^27^ and/or highly expressed in ovules (Extended Data Fig.3h). We found that REM13 exhibited a localization pattern similar to VDD and VAL, demonstrating strong signals over the same set of *clsy3 clsy4*-dependent siRNA loci in Group 1 (Fig. 1e-f and Extended Data Fig.3a-f). For REM19, 80% of its ChIP-seq signals were enriched at Group 1 *clsy3 clsy4*-dependent siRNA loci, with a relatively lower intensity, while 20% extended to Group 2 regions (Fig 1e-f and Extended Data Fig.3a-e). In contrast, 86% of REM8 ChIP-seq peaks localized to Group 2 of *clsy3 clsy4*-dependent siRNA loci instead, showing a preference for *gde1-1* upregulated siRNA regions (Fig 1e-f and Extended Data Fig.3a-c and S3i). In summary, REM transcription factors VDD, VAL, REM12, REM13 and REM19, along with REM22 and REM8 to a lesser extent, were enriched at Group 1 *clsy3 clsy4*-dependent siRNA loci, suggesting functional collaboration in targeting these loci. On the other hand, REM8, together with REM19, localized to a subset of Group 2 of *clsy3 clsy4*-dependent siRNA loci, where the CLSY3/4-Pol IV complex became enriched in the absence of *GDE1*. Collectively, the transcription factors described here can bind over 28% of *clsy3 clsy4*-dependent siRNA loci (Fig. 1h).

## GDE1 controls 24nt-siRNA production and DNA methylation at siren sites in ovules

CLSY3/4 are required for siRNA production at siren sites, highly expressed small RNA producing loci in ovules^10^. Notably, 86 out of 133 siren loci contained the same binding motif as the *clsy3 clsy4* motif1 (Extended Data Fig.4a). REM transcription factor binding sites, including VDD, REM12 and REM13 displayed a similar motif (Extended Data Fig.3f). Overlapping ChIP-seq signals of VDD, VAL and REM13 were observed at 70% of siren loci (Fig. 2a and Extended Data Fig.4b). Additionally, REM19, REM22 and REM8 were enriched at siren loci with somewhat lower enrichment (Extended Data Fig.4b). Furthermore, GDE1 and the Pol IV recruiters, CLSY3 and CLSY4, displayed significant enrichment at siren loci (Fig. 2b and Extended Data Fig.4c). These results suggest that REM-GDE1 complexes may recruit CLSY3/4 to siren loci to regulate siRNA levels.

**Fig. 2.**
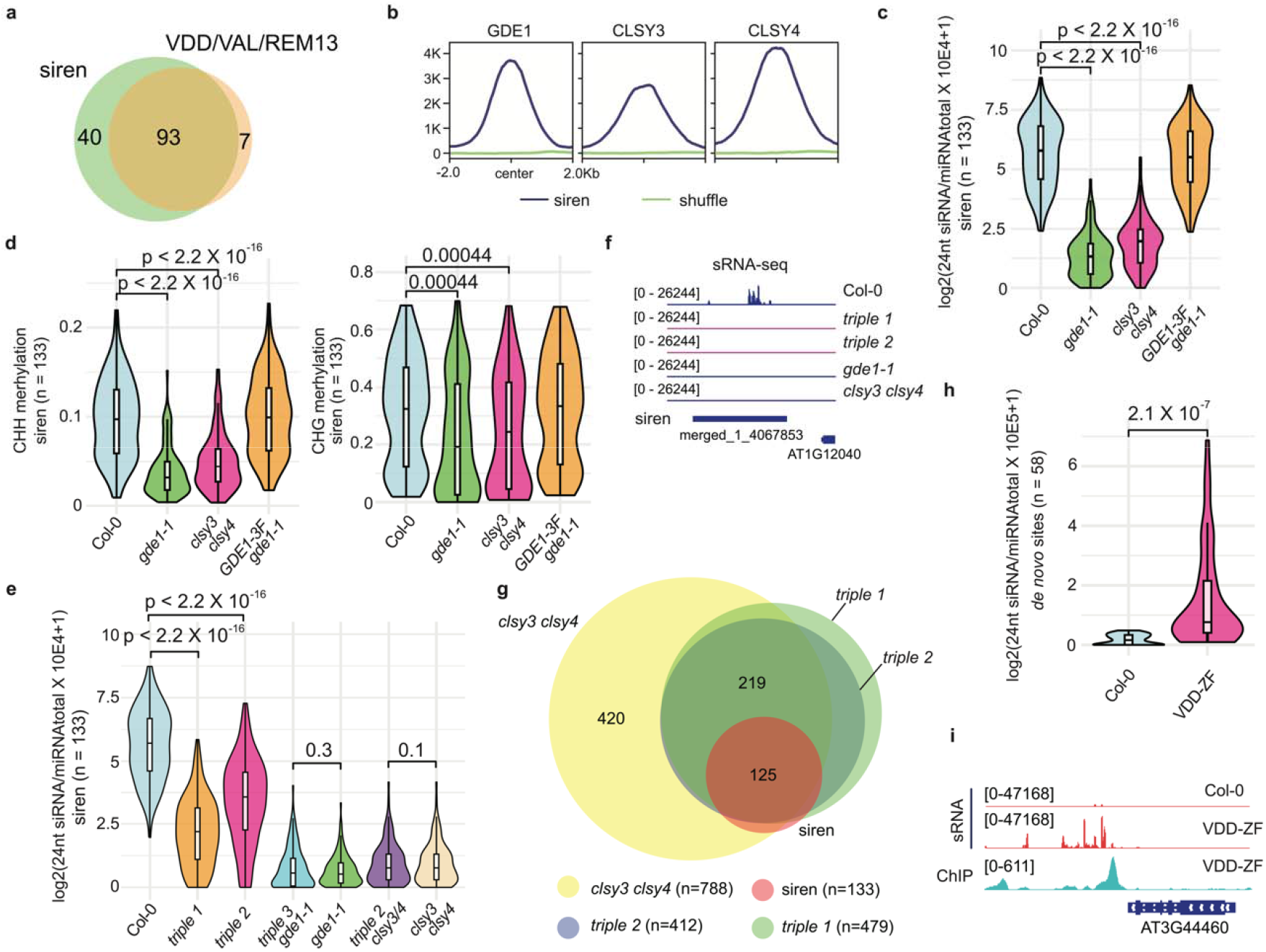
VDD/VAL/REM13-GDE1-CLSY3/4 complex localizes to siren loci for siRNA production. **a**, Venn diagram showing the relationship between siren loci and VDD/VAL/REM13 common ChIP-seq targets. **b**, Metaplot showing enrichment of GDE1, and CLSY3/4 ChIP-seq signal over siren loci. **c**, Violin plot showing the 24nt-siRNA levels at siren loci (n=133) in the indicated genotypes. **d**, Violin plot showing the CHH and CHG methylation levels at siren loci (n=133) in the indicated genotypes. **e**, Violin plot showing the 24nt-siRNA levels at siren loci (n=133) in the indicated genotypes. **f**, Screenshot of siRNA level at a representative siren site. **g**, Venn diagram showing the relationship of down-regulated siRNA loc in each background with siren loci. **h**, Violin plot showing the 24nt-siRNA levels at *de novo* ZF fusion targeted sites (n=58) in the indicated genotypes. **i**, Screenshot of 24nt-siRNA and VDD-ZF ChIP-seq at a *de novo* ZF fusion loci.

The levels of 24nt siRNA at siren loci in *gde1-1* ovules exhibited a strong reduction, phenocopying the *clsy3 clsy4* double mutant (Fig. 2c and Extended Data Fig.4d). This dramatic loss of 24nt siRNA is likely due to the significant loss of CLSY3/4 and Pol IV at siren loci in *gde1-1*, which did not occur at non-siren loci (Extended Data Fig.4e-g). Consistent with the fact that CHH and CHG methylation levels are affected in siRNA biogenesis mutants^28^, we found that these types of methylation were significantly reduced at siren loci in *gde1-1* ovules, resembling the pattern observed in the *clsy3 clsy4* double mutant (Fig. 2d and Extended Data Fig.4h). Taken together, these results indicate that GDE1 is required for CLSY3/4 and Pol IV recruitment, siRNA production, and DNA methylation at siren loci in ovules.

## REM transcription factors promote the biogenesis of 24nt-siRNA

VDD has previously been implicated in controlling the death of the receptive synergid cell during fertilization^25^. Loss-of-function mutants exhibit strong female gametophytic defects^25^. The genetic lethality of knock-out mutants and the multitude of these REM transcription factors make mutant analysis challenging. VAL, REM12 and a VDD interactor, REM46, form a gene cluster in the genome. To create partial loss-of-function mutants of siren targeting REM transcription factors, we utilized the CRISPR/Cas9 editing system to simultaneously delete all three genes. 24nt-siRNA levels at siren loci were significantly reduced in two independent *rem46 val rem12 triple* mutants (Fig. 2e-f and Extended Data Fig.5a), indicating the involvement of these REM transcription factors in regulating siRNA levels at siren loci. Furthermore, the siRNA level at siren loci in *rem46 val rem12 gde1-1* or *rem46 val rem12 clsy3 clsy4* mutants resembled those of their respective parents *gde1-1* or *clsy3 clsy4*, further suggesting that REM transcription factors function in the same pathway with GDE1 and CLSY3/4 at siren loci (Fig. 2e). Additionally, siRNA levels of around 30% of *clsy3 clsy4*-dependent siRNA loci showed reduction in the *rem46 val rem12* mutants although the reduction was not as strong as in the *clsy3 clsy4* mutant (Fig. 2g and Extended Data Fig.5b). All of these reduced *clsy3 clsy4*-dependent siRNA in the *rem46 val rem12* mutants belonged to Group 1. These results indicate that these REM transcription factors regulate siRNA levels at *clsy3 clsy4*-dependent loci.

We also utilized a gain-of-function approach by fusing the REM transcription factor VDD with the artificial zinc finger 108 (ZF), which target hundreds of binding sites throughout the Arabidopsis genome^29,30^. A VDD-ZF fusion was transformed into the WT background and ChIP-seq was performed to identify the binding sites. We identified 397 clear peaks with ZF binding to the genome which were highly enriched for the known ZF binding motif (Extended Data Fig.5c). Small RNA sequencing analysis of the VDD-ZF transgenic plants revealed that 24nt-siRNAs were produced *de novo* at many of the ZF fusion targeted regions in ovule tissues (Fig 2h-i), indicating the sufficiency of REM transcription factors to promote siRNA biogenesis.

To investigate whether the localization of REM transcription factors was affected by GDE1 or CLSY3/4. We performed ChIP-seq using flower tissues collected from VDD-9myc *gde1-1* and VDD-3FLAG *clsy3 clsy4*. We found that the ChIP-seq signals of VDD were reduced significantly at siren loci in both *gde1-1* and *clsy3 clsy4* mutants (Extended Data Fig.5d). These results suggest that GDE1 and CLSY3/4 are both involved in the stabilization of REM transcription factors to siren loci.

## REM-GDE1 complexes recognize the *clsy3 clsy4* motif1 and recruit CLSY3/4-Pol IV to promote unidirectional transcription

The *clsy3 clsy4* motif1 exhibits two or three tandem TTTTGCTTAT sequences with a single nucleotide spaced between them (Fig. 1a and Extended Data Fig.3f). The REM proteins belong to the B3 DNA-binding domain superfamily, and some family members, the ARFs (auxin response factors), possess the ability to form dimers which recognize sequences with a spacer of a specific length^31,32^. Given the fact that REM transcription factors interact with each other *in vivo*, we speculated that each repeat may be bound by a single REM transcription factor. Throughout the genome, the occurrence of the three repeats is rare (n=38), with 48 loci containing two repeats and 37,456 loci having a single repeat. ChIP-seq signals for all components (VDD, VAL, REM13, GDE1, CLSY3, CLSY4, Pol IV) showed strong enrichment at all triple-repeat sites and most double-repeats sites, but very minor enrichment at the single-repeat sites (Extended Data Fig.6a-b). These findings suggest that at least double repeats are required for robust recruitment of the REM transcription factors-GDE1-CLSY3/4-Pol IV transcription complex.

AlphaFold3^33^ confidently predicted that VDD and VAL can dimerize through their N-terminal B3 domain and that each C-terminal B3 domains can bind a repeat in the DNA motif (Fig. 3a and Extended Data Fig.6c). An α-helix of GDE1 (amino acids 249-261) is predicted to insert into a pocket formed by the VDD-VAL dimer to form extensive electrostatic and hydrophobic interactions, and a GDE1 beta sheet (amino acids 604-618) is predicted to be clamped by the C-terminal B3 domains of VDD and VAL (Extended Data Fig.6d). Additionally, a C-terminal α-helix bundle of GDE1 is predicted to form hydrogen-bonds with the C-terminus of the VAL B3 domain (Extended Data Fig.6d). These predictions suggest that VDD and VAL cooperate with GDE1 to recognize the DNA motif, consistent with our *in vivo* results showing a loss of REM binding to the genome in the *gde1-1* mutant.

**Fig. 3.**
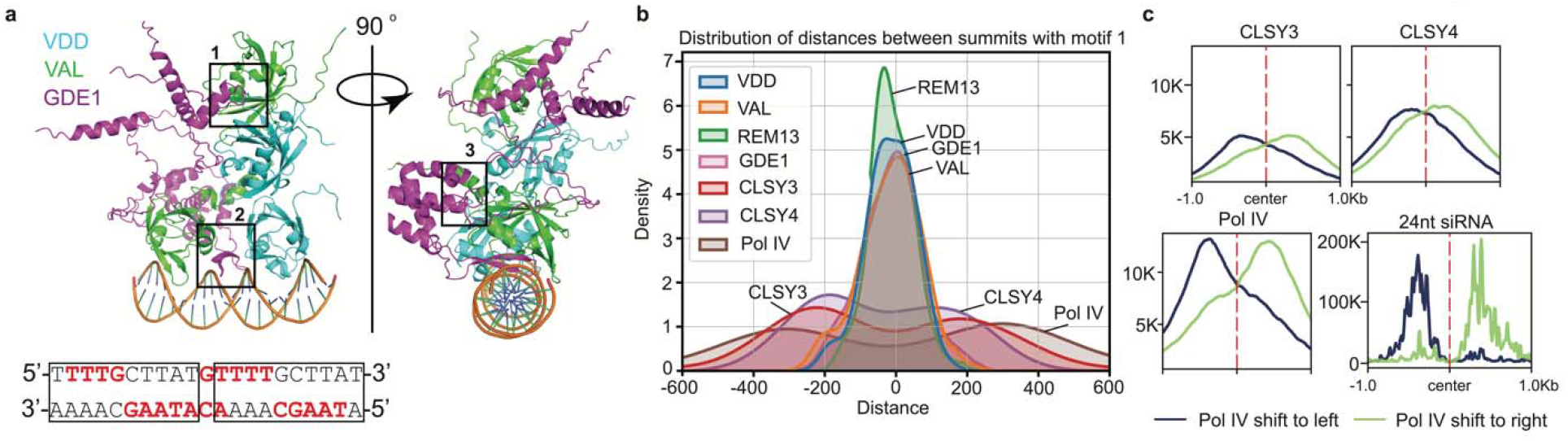
Pol IV transcription complex recognizes *clsy3 clsy4* motif1 and engages in a unidirectional transcription. **a**, AlphaFold3 predicted the structure of VDD-VAL-GDE1 bound to dsDNA containing TTTTGCTTATGTTTTGCTTAT (sequence below, high-affinity residues red bold, two repeats boxed). The DNA is shown as a ribbon representation. **b**, Position of VDD, VAL, REM13, GDE1, CLSY3, CLSY4 and Pol IV ChIP-seq summits to the closest *clsy3 clsy4* motif1 center. **c**, Metaplot showing enrichment of CLSY3, CLSY4, Pol IV ChIP-seq signal and 24nt-siRNA signal over Pol IV left shift (n=23) or Pol IV right shift (n=27).

It was notable that there was always a one-nucleotide space between repeats with strong binding, leading us to speculate that this space might allow for proper spatial dimerization of transcription factors. To examine this further, we identified 27 loci in the genome that contained double repeats but with a two-nucleotide space. Interestingly, we found that no factors in the Pol IV transcription complex were enriched at these loci (Extended Data Fig.7a), underscoring the functional significance of the single space. Consistent with this lack of *in vivo* binding, AlphaFold3 predicted that in the presence of a two-nucleotide spacer, VDD and VAL bound to different sides of the same repeat, and GDE1 failed to orient VDD and VAL properly in order to bind adjacent repeats (Extended Data Fig.7b).

We found that REM transcription factors and GDE1 ChIP-seq peak summits, but not CLSY3/4-Pol IV summits localized to the center of motifs (Fig. 3b and Extended Data Fig.6a). On average, CLSY3/4 and Pol IV were located approximately 200 or 300 base pairs away from the center of the motif, respectively (Fig. 3b). Additionally, CLSY3/4 and Pol IV summits were found on either one side, or the other side, of the center of motif, but not on both sides (Fig. 3c and Extended Data Fig.6a). To further characterize the direction of CLSY3/4/Pol IV distribution relative to the sequence motif, we separated motifs into two groups based on the DNA strand that contained the TTTTGCTTATNTTTTGCTTAT motif sequence in the genome. We found that the distribution of CLSY3/4/Pol IV signals was still localized to either one side or the other side of the motif centers (Extended Data Fig.7c), indicating that other factors, and not the direction of the motif itself, determine the direction of Pol IV transcription. These results suggest that CLSL3/4/Pol IV are recruited to the *clsy3 clsy4* motif1 by the REM transcription factor-GDE1 complex, after which they engage in unidirectional transcription (to one side of the motif or the other) to produce Pol IV transcripts required to produce siRNAs. Consistent with this model, the accumulation levels of 24nt-siRNA transcripts aligned with the localization pattern of the CLSY3/4-Pol IV complex at the *clsy3 clsy4* motif1 (Fig. 3c and Extended Data Fig.7d).

## REM8 recognizes *clsy3 clsy4* motif2 for siRNA production and hyperaccumulates CLSY3/4 to Group 2 sites of *clsy3 clsy4*-dependent siRNA loci in the absence of GDE1

As opposed to the above-described REM transcription factors which appear to mainly drive CLSY3/4-Pol IV to Group 1 of *clsy3 clsy4*-dependent siRNA loci, we found that REM8 exhibited unique binding sites within Group 2 sites (Fig. 1e-f and Extended Data Fig.3a-c and 3i). In line with upregulation of 24nt siRNA in *gde1-1* at Group 2 loci (Fig. 1b), we found that more 24nt siRNAs were detectable at REM8 binding non-siren regions in *gde1-1* (Fig. 4a), which is consistent with the higher CLSY3/4 and Pol IV enrichment at REM8 binding non-siren sites in *gde1-1* background (Extended Data Fig.8a). This also shows that siRNAs accumulate at these sites in a GDE1-independent manner, suggesting that REM8 acts via a different mechanism to recruit Pol IV complexes.

**Fig. 4.**
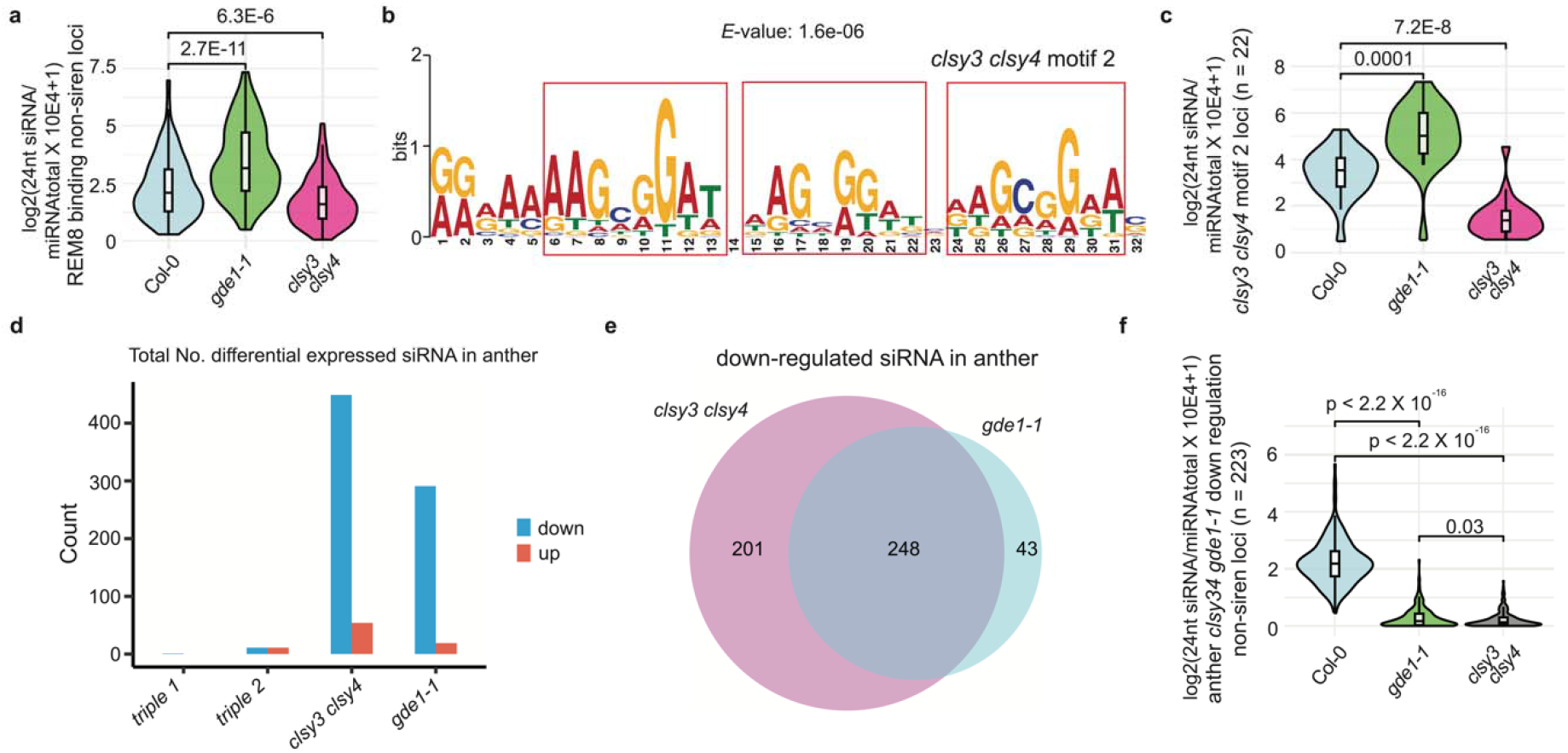
REM8 redistributes Pol IV complex to clsy3 clsy4 motif2 in absence of GDE1 and GDE1 is required for siRNA in anther. **a**, Violin plot showing the 24nt-siRNA levels at REM8 binding non-siren loci (n=162) in the indicated genotypes. **b**, MEME analysis of REM8 binding non-siren loci. **c**, Violin plot showing the 24nt-siRNA levels at *clsy3 clsy4* motif 2 (n=22) in the indicated genotypes. **d**, Total number of differential expressed siRNA in the anther tissue of indicated genotype. **e**, Venn diagram showing the relationship between *clsy3 clsy4*-dependent and *gde1-1*-dependent siRNA loci in anther. **f**, Violin plot showing the 24nt-siRNA levels at anther *clsy3 clsy4* and *gde1-1* down regulation non-siren loci (n=223) in the indicated genotypes.

Motif analysis of REM8 peaks revealed three repeats of the AAGCGGAT sequence with one-nucleotide spaced between them (Fig. 4b) (hereafter named the *clsy3 clsy4* motif 2). 50 *clsy3 clsy4* motif2 were identified, matching to 22 24nt-siRNA loci throughout the genome and REM8 and CLSY3/4 were enriched at most of them (Extended Data Fig.8b). 24nt-siRNA levels were reduced at *clsy3 clsy4* motif2 regions in the *clsy3 clsy4* mutant but increased in the *gde1-1* mutant (Fig. 4c). These results suggest that REM8 can recruit CLSY3/4 to *clsy3 clsy4* motif2 for 24nt-siRNA production, and that this recruitment is more robust in the *gde1-1* mutant, where CLSY3/4-Pol IV are released from the siren loci.

## GDE1 is required for anther siRNA production

CLSY3 is also required for the biogenesis of siRNAs at hyperTE loci in male meiocyte cells^12^. Notably, there is a limited overlap of only 12 shared loci between the maternal siren loci and paternal hyperTE loci^14^, suggesting distinct siRNA biogenesis mechanisms. In tapetum cells, most REM transcription factors, with the exception of REM46 and REM16, are very lowly expressed (Extended Data Fig.3h). Intriguingly, GDE1 stands out as showing very high expression levels in tapetum cells (Extended Data Fig.3h), suggestive of a potential role in meiocyte cells. 449 siRNA loci showed reduction in the anthers of the *clsy3 clsy4* mutant (Fig. 4d), and over half of these loci exhibited similar levels of reduction in the anthers of the *gde1-1* mutant (Fig. 4e). However, hardly any differentially expressed siRNAs were found in *rem46 val rem12 triple* mutants (Fig. 4d). Consistently, only GDE1, CLSY3/4, and Pol IV, but none of the tested REM transcription factors showed enrichment at anther *clsy3 clsy4*-dependent siRNA loci (Extended Data Fig.9). Together, these results suggest that GDE1 is also required for anther siRNA production, but likely via a different mechanism than that seen in ovules.

## Conclusion

It has been previously observed that CLSYs control all Pol IV-dependent 24nt-siRNA production, with CLSY1/2 associating with SHH1, an H3K9 methylation reader to recruit Pol IV transcription complexes to regulate 24nt-siRNA production^9,34-37^. In this way, histone marks (other epigenetic modifications) are influencing Pol IV transcription. In contrast, our study reveals that CLSY3/4 directs Pol IV binding and siRNA production in part via the novel factor GDE1, which acts in concert with REM transcription factors to direct siRNA production to specific sequences in ovules. This work suggests that CLSY proteins have evolved multiple distinct mechanisms, some based on epigenetic information, and some based on genetic information, for recruiting Pol IV to specific sites.

Transcription factors are known to assist in the recruitment of DNA methylation and gene silencing in mammals. For example, KRAB-zinc finger proteins (KRAB-ZFPs), the largest transcription factor family in mammalian cells, are known to bind specific DNA sequences and recruit the repressor KRAB-associated protein 1 (KAP-1)^38,39^ which recruits diverse complexes to regulate DNA methylation, histone deacetylation, H3K9 trimethylation and transposon and gene silencing^40,41^. In an analogous fashion, our results suggest that REM transcription factors recruit CLSY3/4-Pol IV complexes with the help of GDE1 to direct siRNA production and subsequent DNA methylation. These parallels suggest that sequence-specific DNA binding proteins are used widely throughout different eukaryotic species to properly pattern DNA methylation.

## Materials and Methods

### Plant materials and growth conditions

All Arabidopsis plants used in this paper are Col-0 ecotype, and plants are grown under standard condition with 16□h light/8□h dark at 22□°C. The T-DNA insertion lines used in this study included *gde1-1* (SALKseq_10069.1), *clsy3-1* (SALK_040366), and *clsy4-1* (SALK_003876). The Agrobacterium (AGL0 strain) mediated floral dipping was used to generate all the transgenic plants.

### Plasmid construction

*pGDE1:GDE1-3FLAG, pCLSY3:CLSY3-9myc, pCLSY4:CLSY4-9myc, pPol IV:Pol IV-9myc, pVDD:VDD-9myc, pVDD:VDD-3FLAG, pVAL:VAL-9myc, pREM13:REM13-9myc, pREM19:REM19-9myc, pREM22:REM22-9myc, pREM8:REM8-9myc, pVDD:VDD-FLAG-*

*ZF108*: the genomic DNA sequences of each with promoter sequences (around 2□kb upstream from the start codon) were first cloned into *pENTR/D-TOPO* vectors (Invitrogen), and then to the destination vector *pEG302-GW-3FLAG, pEG302-GW-9Myc*, or *pEG302-GW-3FLAG-ZF108* by LR reaction (LR Clonase II, Invitrogen).

### Chromatin immunoprecipitation sequencing

Fifteen ml of floral tissues were used for each ChIP. The plant materials were ground into fine powder with liquid nitrogen and resuspended with nuclei isolation buffer (50□mM HEPES, 1□M sucrose, 5□mM KCl, 5□mM MgCl_2_, 0.6% Triton X-100, 0.4□mM PMSF, 5□mM benzamidine, 1% formaldehyde (Sigma), and 1X Protease Inhibitor Cocktail (Roche)) for 10□min with rotation. 1.7□ml of 2□M glycine solution was added immediately to stop the crosslinking. Lysates were filtered through Miracloth and the nuclei were collected by centrifuge at 4□°C with 2880×g for 20□min. The pellet was resuspended in 1 ml of extraction buffer 2 (0.25□M sucrose, 10□mM Tris-HCl pH 8.0, 10□mM MgCl_2_, 1% Triton X-100, 5□mM BME, 0.1□mM PMSF, 5□mM Benzamidine, and 1x Protease Inhibitor Cocktail (Roche)) and centrifuged at 12,000 x g at 4□°C for 10□minutes. The nuclei were then resuspended with extraction buffer 3 (1.7□M sucrose, 10□mM Tris-HCl pH 8.0, 2□mM MgCl_2_, 0.15% Triton X-100, 5□mM BME, 0.1□mM PMSF, 5□mM Benzamidine, and 1x Protease Inhibitor Cocktail (Roche)), at 4□°C with 12,000×g for 60□min. The relative pure nuclei were lysed with 400□µL nucleic lysis buffer (50□mM Tris-HCl pH 8.0, 10□mM EDTA, 1% SDS, 0.1□mM PMSF, 5□mM Benzamidine, and 1x Protease Inhibitor Cocktail (Roche)) on ice for 10□minutes and a total of 1.7□ml of ChIP Dilution Buffer (1.1% Triton X-100, 1.2□mM EDTA, 16.7□mM Tris pH 8.0, 167□mM NaCl, 0.1□mM PMSF, 5□mM Benzamidine, 1x protease inhibitor cocktail tablet) was added to the lysed nuclei. Chromatin was sheared by Bioruptor Plus (Diagenode) for 30 cycles with 30□s on/30□s off per cycle. The lysate was centrifuged twice at 4□°C with 20,000×g for 10□min, and the supernatant was incubated with either FLAG epitope (Sigma F1804) or myc epitope (Cell Signaling, 71D10) at 4□°C overnight. Next, the magnetic Protein A and Protein G Dynabeads (Invitrogen) were added and inoculated at 4□°C for 2□h with rotation. The beads were washed with low salt solution twice (150□mM NaCl, 0.2% SDS, 0.5% Triton x-100, 2□mM EDTA, and Tris pH 8.0), high salt solution (150□mM NaCl, 0.2% SDS, 0.5% Triton x-100, 2□mM EDTA, and Tris pH 8.0), LiCl solution (250□mM LiCl, 1% Igepal, 1% Sodium Deoxycholate, 1□mM EDTA, and 10□mM Tris pH 8.0), and TE solution (1□mM EDTA and 10□mM Tris pH 8.0) for 5□minutes at 4□°C, respectively. The chromatin was eluted with elution buffer (1% SDS, 10□mM EDTA, and 0.1□M NaHCO_3_), and subjected to reverse crosslinking by adding 20□µL 5□M NaCl and incubated at 65□°C overnight. Then 1□μL Protease K (20□mg/mL, Invitrogen), 10□μL of 0.5□M EDTA pH 8.0, and 20□μL 1□M Tris pH 6.5 were added to deactivate the protein for 4□hours at 45□°C and DNA was purified through phase lock gel (VWR) and precipitated with 1/10 volume of 3□M Sodium Acetate (Invitrogen), 2□μL GlycoBlue (Invitrogen), and 1□mL 100% Ethanol at −20□°C overnight. The precipitated DNA was used for library construction following the manual of the Ovation Ultra Low System V2 kit (NuGEN), and the libraries were sequenced on Illumina NovaSeq 6000 or HiSeq 4000 instruments.

### IP-MS

50□mL of floral tissues from FLAG epitope tagged transgenic plants were used for each IP-MS experiment and floral tissues of Col-0 plants were used as control. Flower tissue was ground to fine powder in liquid nitrogen with homogenizer. Tissue powder was completely resuspended in 25□mL IP buffer (50□mM Tris-HCl pH 8.0, 150□mM NaCl, 5□mM EDTA, 10% glycerol, 0.1% Tergitol, 0.5□mM DTT, 1□mg/mL Pepstatin A, 1□mM PMSF, 50□µM MG132 and cOmplete EDTA-free Protease Inhibitor Cocktail (Roche)) at 4□°C for 10□minutes with rotation. The issue was further disrupted with dounce homogenizer. The lysates were filtered with Miracloth and centrifuged at 20,000□g for 10□min at 4□°C. The supernatant was incubated with 250□μL anti-FLAG M2 magnetic beads (Sigma) at 4□°C for 2□h with rotation. The magnetic beads were washed 4 times with IP buffer and eluted with 250□µg/mL 3xFLAG peptides. Eluted proteins were used for Trichloroacetic acid (TCA) precipitation and mass spectrometric analysis.

### Quantitative proteomics

Protein pellets were resuspended in 8□M urea and 100□mM Tris pH□8.5, then reduced by adding Tris (2-carboxyethyl) phosphine (TCEP) to a final concentration of 5□mM and incubation for 30□min. Next, the proteins were alkylated by adding iodoacetamide to a final concentration of 10□mM for another 30□min at room temperature. Before protein digestion, the urea concentration was diluted to 2□M with 100□mM Tris pH□8.5. Then, the proteins were digested with LysC (BioLabs) at a 1:100 enzyme/protein ratio at 37□°C for 4□hours, followed by the trypsin digestion at 1:100 (trypsin: protein) at 37□°C for 12□h. To stop the digestion, 5% formic acid was added to the samples. Next, the peptides were desalted using C18 pipette tips (Thermo Scientific) and reconstituted in 5% formic acid before being analyzed by LC-MS/MS. Tryptic peptide mixtures were loaded onto a 25□cm long, 75□μm inner diameter fused-silica capillary, packed in-house with bulk 1.9□μM ReproSil-Pur beads with 120□Å pores as described(*1*). The peptides were delivered by a 140-min water-acetonitrile linear gradient in 6–28% buffer (acetonitrile solution, 0.1% formic acid and 3% DMSO) using a Dionex Ultimate 3000 nanoflow UHPLC (Thermo Scientific), at a flow rate of 200 nL/min, further increased to 35% and followed by a rapid ramp-up to 85%. The eluted peptides were ionized and the Orbitrap Fusion Lumos Tribrid Mass Spectrometer (Thermo Scientific) was used to acquire the Mass Spectrometer. The data-dependent acquisition strategy consisted of a repeating cycle of a full MS1 spectrum (Resolution = 120,000) followed by sequential MS2 scan (Resolution = 15,000). Label-free quantification was performed using the MaxQuant software package (v1.6.17.0) with LFQ default setting(*2*), and Arabidopsis TAIR 10 proteome database was used for the database search. Trypsin digestion was applied and a maximum of two missed cleavages were allowed in all searches for tryptic peptides of length 8–40 amino acids. In all, 1% false discovery rate was used as a filter at both protein and peptide-spectrum match (PSM) levels. IP-MS of Col-0 plant tissue was used as control. The empirical Bayes test performed by LIMMA was used for statistical analysis.

### Co-immunoprecipitation

10 ml of floral tissues was collected from GDE1-3FLAG x VDD-9myc, GDE1-3FLAG x CLSY3-9myc, GDE1-3FLAG x Pol IV-9myc, VDD-9myc, CLSY3-9myc, and Pol IV-myc. Tissues were ground into fine powder with liquid nitrogen and mixed with 10□mL IP buffer (50□mM Tris HCl pH 7.5, 150□mM NaCl, 2□mM EDTA, 2□mm DTT, 0.8% TritonX-100, and 1x Protease inhibitor (Roche)), and incubated at 4□°C for 20□min. The lysate was centrifuged at 18,000×g for 10□min at 4□°C, and the supernatant was centrifuged one more time. The supernatant was incubated with 30□μL anti-FLAG M2 magnetic beads (Sigma) for 2□h at 4□°C. The beads were washed with IP buffer for 5 times, and proteins were eluted with 40□μL elution buffer (IP buffer containing 100□μg/mL 3xFLAG peptide as final concentration) by vigorously shaking at 37□°C for 15 mins. The elutions were mixed with 2x SDS loading buffer for western blot.

### Small RNA-seq

Pistils and anther (stage 9 or yonger) of each genotype were collected and ground into fine powder with liquid nitrogen. Total RNA was extracted using the Direct-zol RNA Miniprep kit (Zymo) according to the manufacturer’s instructions. Two micrograms of total RNA were mixed with equal volume of the 2x RNA loading dye (*3*), denatured at 65°C for 10 min and immediately chilled on ice. Denatured total RNA was separated on 15% TBE urea gel (Invitrogen) and small RNAs from 15-30 nts were excised. The excised gel pieces were mashed by pestles. Small RNA was eluted with 400 ul nuclease-free water at 70°C for 10 mins and then precipitated with ethanol. Small RNA libraries were made using NEBNext® Small RNA Library Prep Set for Illumina® (Multiplex Compatible) (*3*) according to the manufacturer’s instructions. The libraries were sequenced on Illumina NovaSeq 6000 or HiSeq 4000 instruments.

### RNA-seq

Seedlings, pistils, and pollens of Col-0 were collected and ground into fine powder with liquid nitrogen. Total RNA was extracted using the Direct-zol RNA Miniprep kit (Zymo) according to the manufacturer’s instructions. One microgram of total RNA was used to prepare the libraries for RNA-seq following TruSeq Stranded mRNA kit (Illumina), and the libraries were sequenced on Illumina NovaSeq 6000 or HiSeq 4000 instruments.

### Whole genome bisulfite sequencing (WGBS)

DNA from pistils were extracted with Qiagen DNeasy plant mini kit (Qiagen 69106). RNA was removed with PureLink RNase A (Invitrogen). A total of 100 ng of DNA was sheared to 200 bp with a Covaris S2 (Covaris). Libraries were prepared with the Epitect Bisulfite Conversion kit (QIAGEN) and the Ovation Ultralow Methyl-seq kit (NuGEN) following the manufacturer’s instructions. The libraries were sequenced on Illumina NovaSeq 6000 or HiSeq 4000 instruments.

### Alphafold3 structure prediction

The 3D models of VDD-VAL-GDE1-motif1 and VDD-VAL-motif1 were generated by Alphafold3(*3*), and visualized and analyzed by PyMOL 3.0.2.

### Bioinformatic analysis

#### ChIP-seq analysis

Raw reads were aligned to the Arabidopsis reference genome (TAIR10) with Bowtie2 (v2.3.4.3)(*4*), allowing only uniquely mapped reads with perfect matches. The Samtools version 1.9 was used to remove duplicated reads(*5*). The deeptools version 3.1.3 was used to generate Bigwig tracks(*6*). Peaks were called using MACS2 (v2.1.1)(*7*).

#### Differentially ChIP-seq localization analysis

ChIP-seq levels at the *clsy3 clsy4*-dependent siRNA regions were quantified with the HOMER annotatePeaks.pl script using the “-noadj, -size given and -len 1” options. Differentially expressed 24nt-siRNA compared to the WT controls were then identified using DESeq (log2 FC□≥□1 and FDR□≤□0.05).

#### Binding motif analysis

MEME 5.5.0 was used to discover the motifs of the ChIP-seq data sets(*8*). FIMO was used to scan genome-wide distributions of *clsy3 clsy4* motif1 TTTTGCTTAT (single-repeat) with one mismatch allowed, TTTTGCTTATNTTTTGCTTAT (double-repeats) with one mismatch allowed in each repeat, TTTTGCTTATNTTTTGCTTATNTTTTGCTTAT (triple-repeats) with one mismatch allowed in each repeat, and TTTTGCTTATNNTTTTGCTTAT (double-repeats with a two-nucleotide space) with one mismatch allowed in each repeat(*9*); *clsy3 clsy4* motif2 AAGCGGATNAAGCGGATNAAGCGGAT with p-value less than 5E-09 and q-value less than 0.025. Tomtom was used to analyze the similarities between motifs(*10*).

#### WGBS analysis

WGBS raw reads were aligned to both strands of reference genome TAIR10 using BSMAP (v.2.74)(*11*) with allowing up to 2 mismatches and 1 best hit (-v 2 -w 1). Reads with more than 3 consecutively methylated CHH sites were considered as non-converted reads and removed. Methylation levels were calculated with the ratio of C/(C + T).

#### RNA-seq analysis

Col-0 leaf, meiocyte and tapetum RNA seq data were downloaded from NCBI Gene Expression Omnibus (GEO) as accession GSM2306324; GSM2306325; GSM2306326(*12*), GSM2306313; GSM2306314; GSM2306315(*12*) and GSM4911399; GSM4911400; GSM4911401(*13*), respectively. All raw reads of RNA-seq data were aligned to reference genome TAIR10 by Bowtie2 (v2.3.4.3)(*4*), and expression abundance was calculated by RSEM (v1.3.1) with default settings(*14*). The bamCoverage of deeptools version 3.1.351 was used to normalize the data with RPKM (*6*).

#### Small RNA-seq analysis

Adaptor sequence (TGGAATTCTCGG) of small RNA-seq reads were trimmed with trim_galore, and trimmed reads were mapped to the reference genome TAIR10 using Bowtie2 with only one unique hit and zero mismatches(*4*). sRNA reads that mapped to chloroplast, mitochondrial DNA, tRNA, rRNA, small nucleolar RNAs (snoRNAs), and small nuclear RNAs (snRNAs) were removed using bedtools (v2.26.0)(*15*). The deeptools version 3.1.3 was used to generate Bigwig tracks(*6*). The bamCoverage of deeptools version 3.1.351 was used to normalize the data with RPKM(*6*).

#### Differentially expressed (DE) 24nt-siRNA clusters analysis

Pol IV dependent master siRNA were defined from a previous publication(*16*). 24nt-siRNA levels at the master 24nt-siRNA were quantified with the HOMER annotatePeaks.pl script using the “-noadj, -size given and -len 1” options. 24nt-siRNA expression levels were normalized by total miRNA amount, which were defined from previously (*17*). Differentially expressed 24nt-siRNA compared to the WT controls were then identified using DESeq (log2 FC□≤□1 and FDR□≤□0.05).

## Acknowledgments

We thank Dr. Julie Law and Dr. Guanghui Xu at Salk Institute for discussion and sharing the *clsy* mutants; and Dr. Jiamu Du at Southern University of Science and Technology for discussion of the Alphafold3 results; Dr. Colette Picard, Dr. Ming Wang, and Dr. Yan He for advice on the manuscript; and Jessica Deng, Yoshiaki Yamamoto, Joowon Um, Carsten Hoeke, and Ziqi Guo for technical support. We also thank Mahnaz Akhavan and the UCLA BSCRC BioSequencing Core for sequencing support.

## Author contributions

ZW and SEJ designed the research, interpreted data, and wrote the manuscript; ZW performed most of the experiments and bioinformatic data analysis; YX initiated the project and generated GDE1 tagged transgenic lines; SW performed siRNA and transcription factors binding DE analysis; YHS generated transcription factors tagged lines and performed ChIP-seq. ZZ performed WGBS bioinformatic data analysis and initial bioinformatic data analysis of sRNA-seq, RNA-seq, ChIP-seq; SF performed high throughput sequencing; ZW, JS, and JW performed IP-MS and interpreted the data. JD and AL provided technical help.

## Competing interests

Authors declare that they have no competing interests.

## Materials & Correspondence

Steven E. Jacobsen

## Data and materials availability

All the high-throughput sequencing data generated in this study is accessible at NCBI’s Gene Expression Omnibus (GEO) via GEO Series accession number.

## Notes

### Competing Interest Statement

The authors have declared no competing interest.

