## Extended data figures for "Arabidopsis REM transcription factors and GDE1 shape the DNA methylation landscape through the recruitment of RNA Polymerase IV transcription complexes"

Extended Data 1

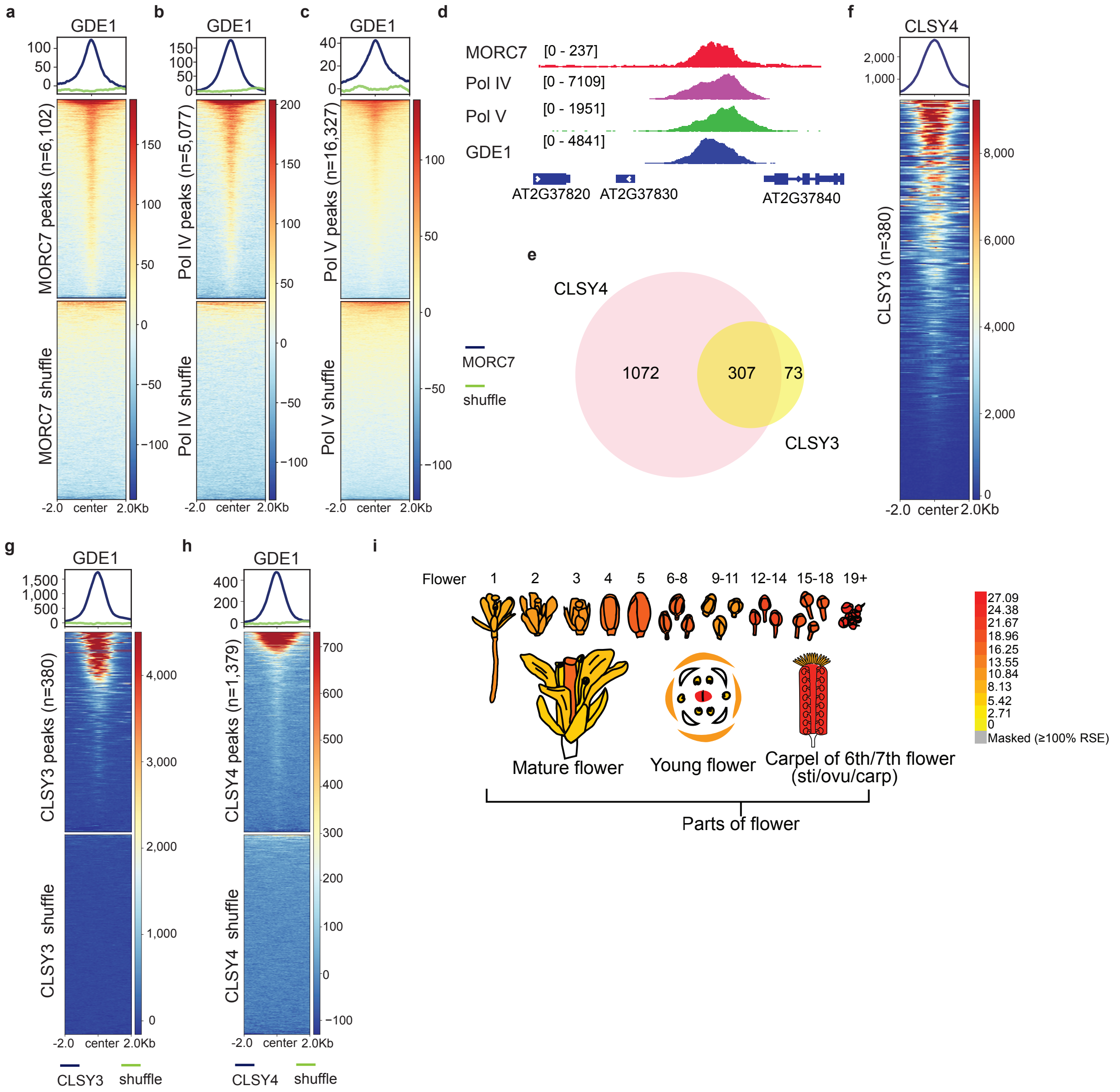


**Extended Data Figure 1 | GDE1 is a RdDM protein and co-localizes with Pol IV recruiter CLSY3/4. a, b, c,** Metaplot and heatmap showing enrichment of GDE1 ChIP-seq signal over MORC7 (n=6102) (**a**), Pol IV (n=5077) (**b**), and Pol V (n=16,327) (**c**). **d,** Screenshot of MORC7, Pol IV, Pol V and GDE1 ChIP-seq at a representative locus. **e,** Venn diagram showing the relationship between CLSY3 and CLSY4 binding sites. **f,** Metaplot and heatmap showing enrichment of CLSY4 ChIP-seq signal over CLSY3 (n=380). **g, h,** Metaplot and heatmap showing enrichment of GDE1 ChIP-seq signal over CLSY3 (n=380) (**g**), and CLSY4 (n=1368) (**h**). **i,** Relative expression levels of the GDE1 genes in select tissues from ePlant expression viewers.

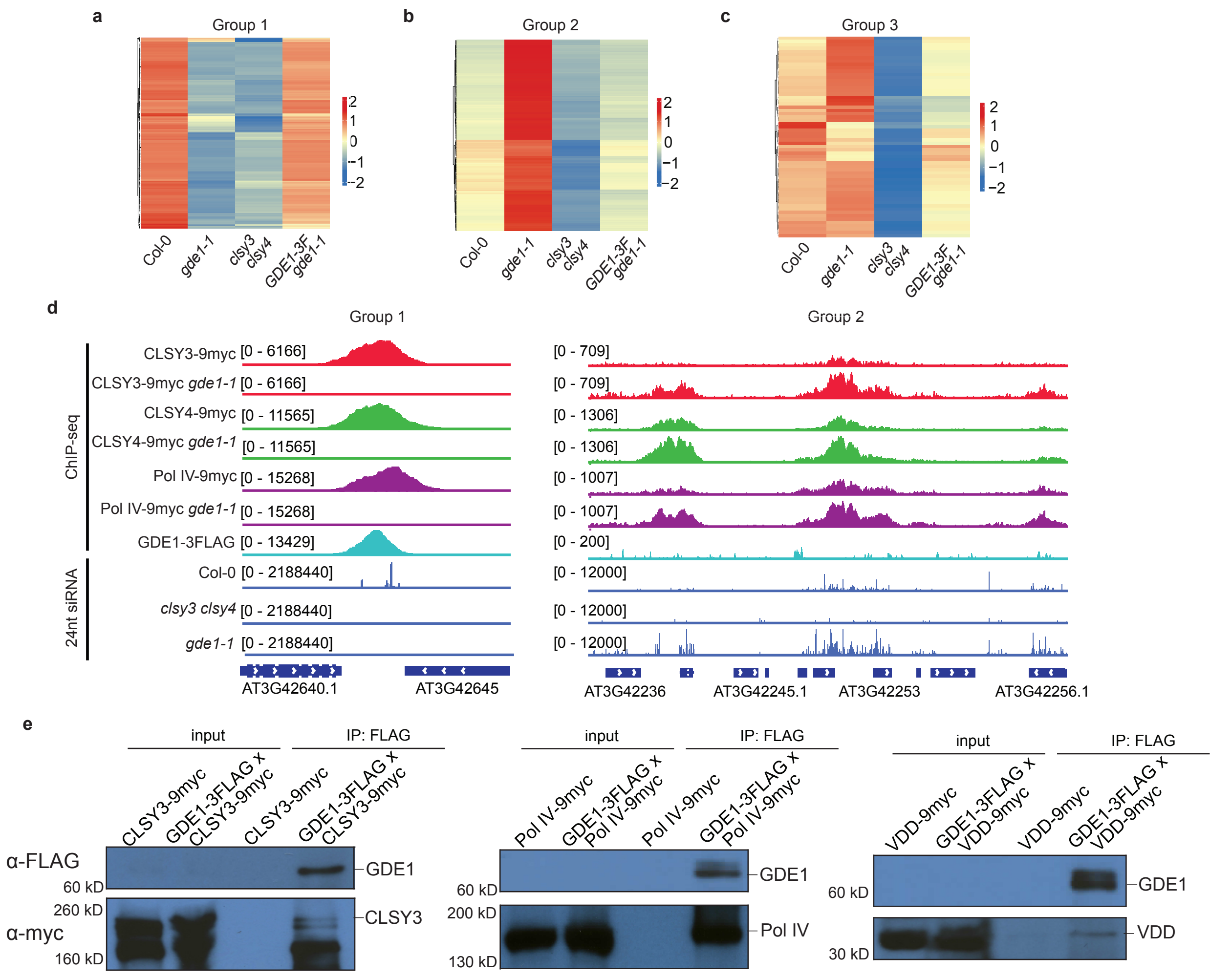


**Extended Data Figure 2 | GDE1 co-localizes and interacts with Pol IV complex and REM transcription factors for siRNA production. a, b, c,** Heatmap showing 24nt-siRNA levels in Group1 (**a**), Group 2 (**b**) or Group3 (**c**) of *clsy3 clsy4*-dependent siRNA sites. **d,** Screenshot of CLSY3-9myc, CLSY3-9myc *gde1-1*, CLSY4-9myc, CLSY4-9myc *gde1-1*, Pol IV-9myc, Pol IV-9myc *gde1-1* and GDE1-3FLAG ChIP-seq signals with control ChIP-seq signals subtracted over a Group1 (*clsy3 clsy4*-dependent, *gde1*-dependent) representative locus (left) or three Group2 (*clsy3 clsy4*-dependent, *gde1*-upregulated) representative loci (right). **e,** Western blot showing a Co-IP assay in *GDE1-3FLAG* F2 crossed lines with *CLSY3-9myc* (left), *Pol IV-9myc* (middle) or *VDD-9myc* (right), respectively. Similar results were observed in two biological replicates.

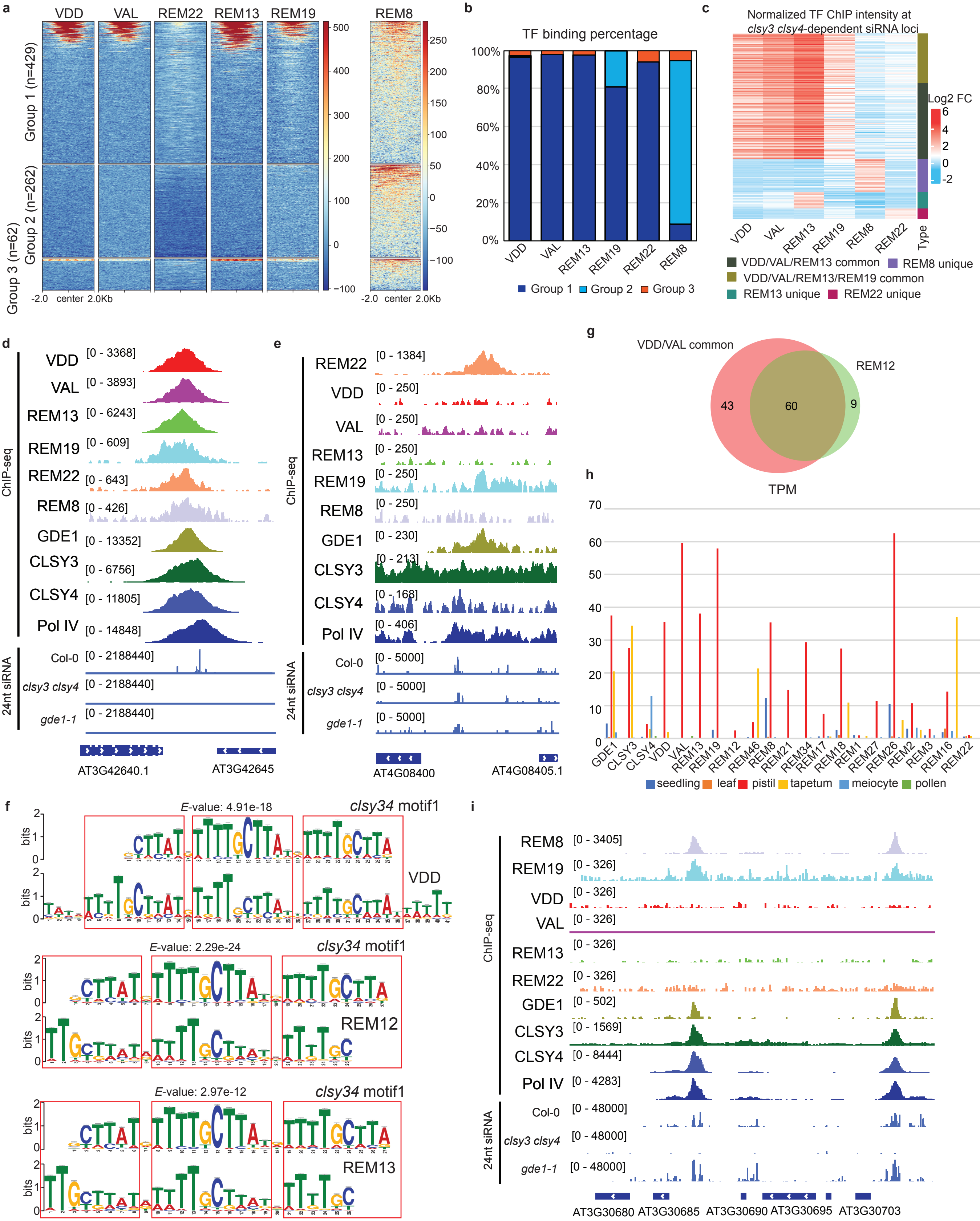
a
VDD
VAL
REM22
REM13
REM19
REM8
Group 1 (n=429)
Group 2 (n=262)
Group 3 (n=62)
-2.0 center 2.0Kb
b
TF binding percentage
Group 1
Group 2
Group 3
c
Normalized TF ChIP intensity at clsy3 clsy4-dependent siRNA loci
Log2 FC
Type
VDD/VAL/REM13 common
REM8 unique
VDD/VAL/REM13/REM19 common
REM13 unique
REM22 unique
d
ChIP-seq
24nt siRNA
GDE1
CLSY3
CLSY4
Pol IV
Col-0
clsy3 clsy4
gde1-1
AT3G42640.1
AT3G42645
e
AT4G08400
AT4G08405.1
g
VDD/VAL common
REM12
43
60
9
h
TPM
seedling
leaf
pistil
tapetum
meiocyte
pollen
f
clsy34 motif1
E-value: 4.91e-18
E-value: 2.29e-24
E-value: 2.97e-12
REM12
REM13
bits
i
AT3G30680
AT3G30685
AT3G30690
AT3G30695
AT3G30703

**Extended Data Figure 3 | Differential localization of REM transcription factors at *clsy3 clsy4*-dependent siRNA sites. a,** Heatmap showing enrichment of VDD, VAL, REM22, REM13, REM19 and REM8 ChIP-seq signal over Group1, Group2, and Group3 of *clsy3 clsy4*-dependent siRNA sites. **b,** Bar plot showing percentage of each REM transcription factors binding to three distinct groups of *clsy3 clsy4*-dependent siRNA loci. **c,** Heatmap showing normalized transcription factors ChIP-seq intensity at *clsy3 clsy4*-dependent siRNA loci. **d,** Screenshot of VDD, VAL, REM13, REM19, REM22, REM8, GDE1, CLSY3, CLSY4 and Pol IV ChIP-seq at a representative Group 1 of *clsy3 clsy4*-dependent siRNA sites. **e,** Screenshot of REM22, VDD, VAL, REM13, REM19, REM8, GDE1, CLSY3, CLSY4 and Pol IV ChIP-seq at a distinct representative Group 1 of *clsy3 clsy4*-dependent siRNA sites, where REM22, but not other REM proteins exhibit enrichment. **f,** TOMTOM analysis showing the similarities of VDD, REM12 and REM13 binding sites with *clsy3 clsy4* motif1. **g,** Venn diagram showing the similarities between VDD/VAL common targeted siRNA loci with REM12 targeted siRNA loci. **h,** RNA-seq analysis of REM transcription factors, GDE1 and CLSY3/4 in different tissues. **i,** Screenshot of REM8, REM19, VDD, VAL, REM13, REM22, GDE1, CLSY3, CLSY4 and Pol IV ChIP-seq at two representative Group 2 of *clsy3 clsy4*-dependent siRNA loci.

### Extended Data 4

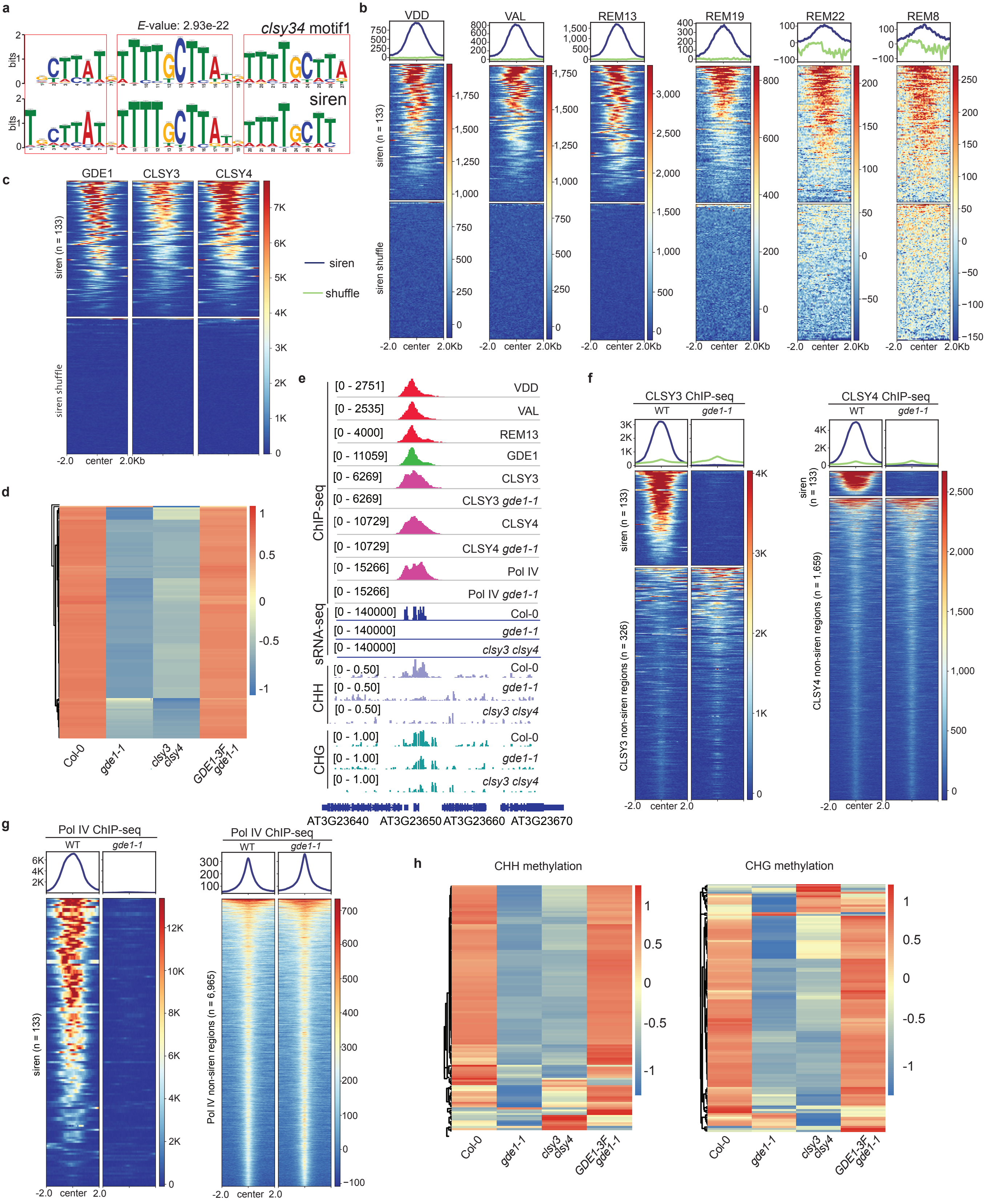

**Extended Data Figure 4 | Transcription factor driven Pol IV transcription complex localizes to siren loci for siRNA production and DNA methylation. a,** TOMTOM analysis showing the similarities of siren loci and *clsy3 clsy4* motif1. **b,** Metaplot and heatmap showing enrichment of VDD, VAL, REM13, REM19, REM22 and REM8 ChIP-seq signal over siren loci (n=133). **c,** Heatmap showing enrichment of GDE1, CLSY3 and CLSY4 ChIP-seq signal over siren loci (n=133). **d,** Heatmap showing the 24nt-siRNA levels at siren loci (n=133) in the indicated genotypes. **e,** Screenshots of VDD, VAL, REM13, GDE1, CLSY3, CLSY3 *gde1-1*, CLSY4, CLSY4 *gde1-1*, Pol IV and Pol IV *gde1-1* ChIP-seq signals with control ChIP-seq signals subtracted and 24nt-siRNA by sRNA-seq and CHG/CHH DNA methylation level by WGBS over a representative siren site in the indicated genotypes. **f,** Metaplot and heatmap showing enrichment of CLSY3/4 ChIP-seq signal over CLSY3/4-bound siren loci (n=133) and CLSY3/4 non-siren regions in WT and *gde1-1* backgrounds. **g,** Metaplot and heatmap showing enrichment of Pol IV ChIP-seq signal over Pol IV-bound siren loci (n=133) and Pol IV non-siren regions in WT and *gde1-1* backgrounds. **h,** Heatmap showing the CHH and CHG methylation levels at siren loci (n=133) in the indicated genotypes.

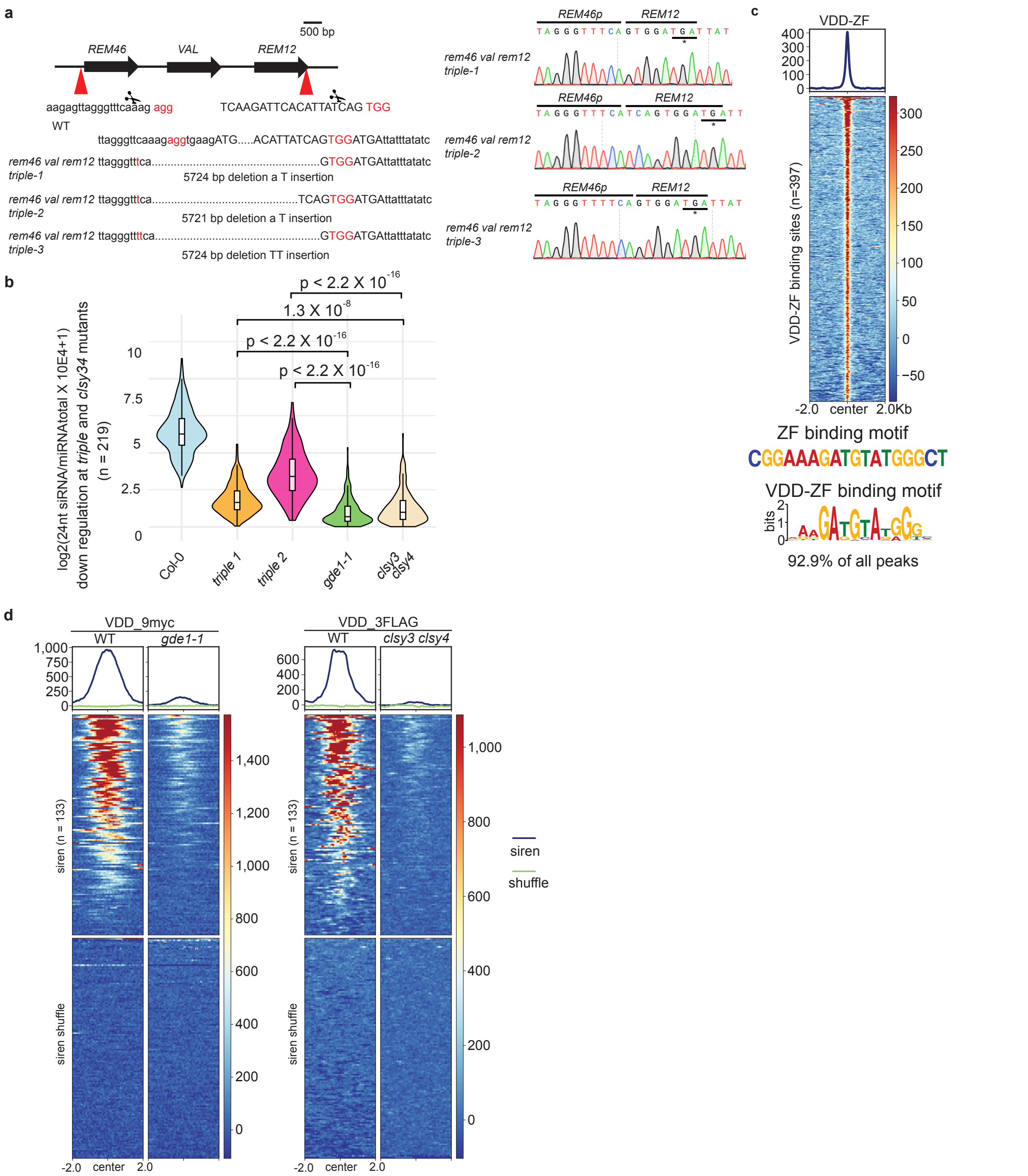

**Extended Data Figure 5 | REM transcription factors are required for siRNA production at *clsy3 clsy4*-dependent siRNA loci. a,** Schematic description of CRISPR/Cas9 construct design for knocking out the tandemly duplicated REM46, VAL, and REM12. Deletion detected in genomic DNA of *triple* mutants are shown with chromatographs from Sanger sequencing. **b,** Violin plot showing the 24nt-siRNA levels at siRNA down regulation loci at both *triple* and *clsy3 clsy4* mutants (n=219) in the indicated genotypes. **c,** Metaplot and heatmap showing enrichment of ZF-VDD ChIP-seq signal over ZF fusion targeted sites (n=397) and motif identified by MEME motif analysis for ZF-VDD. **d,** Metaplot and heatmap showing enrichment of VDD ChIP-seq signal over siren loci in Col-0, *gde1-1* and *clsy3 clsy4* double backgrounds.

Extended Data 6

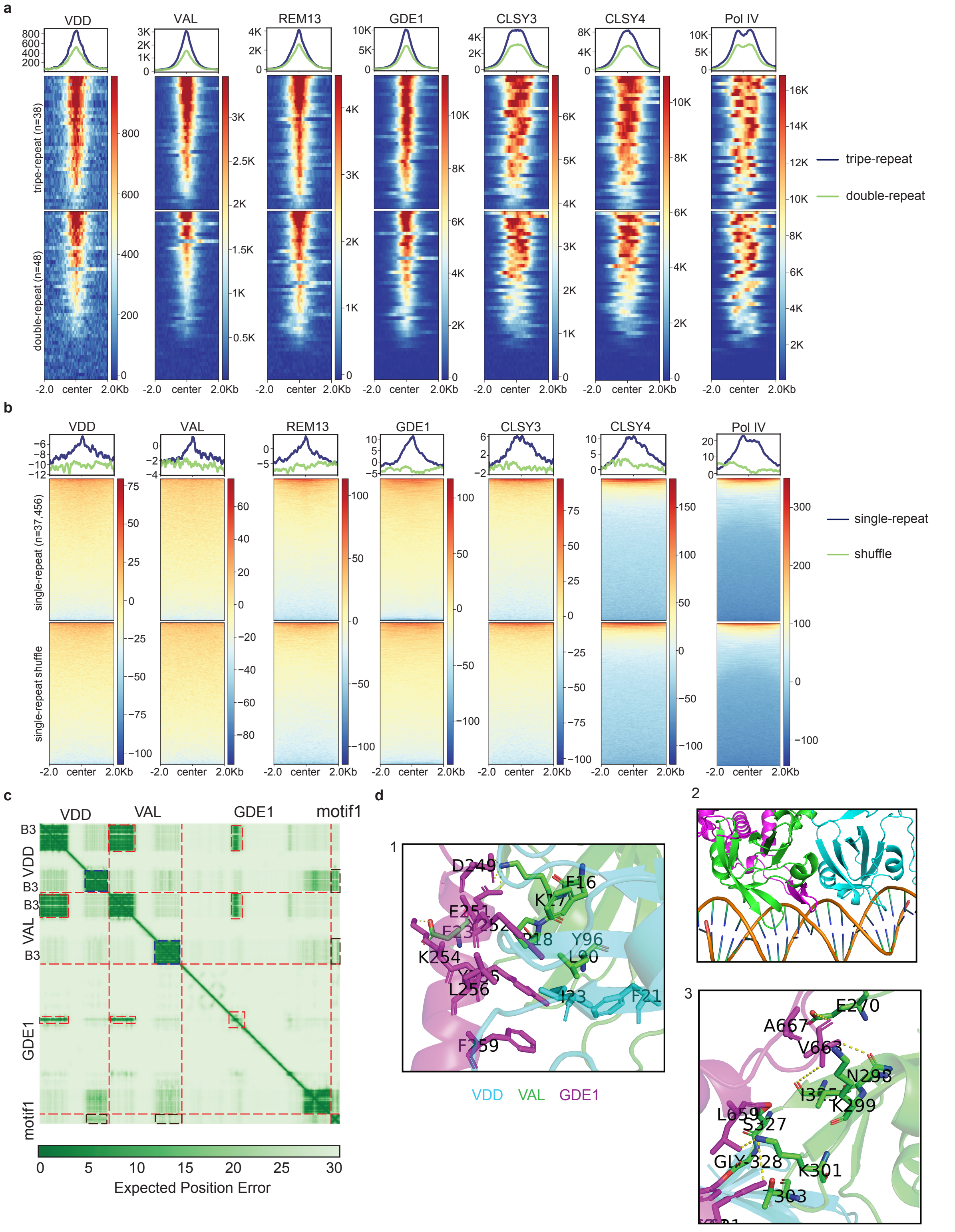


**Extended Data Figure 6 | Pol IV transcription complex recognizes *clsy3 clsy4* motif1. a,** Metaplot and heatmap showing enrichment of VDD, VAL, REM13, GDE1, CLSY3, CLSY4 and Pol IV ChIP-seq signal over tripe-repeats sites (n=38) and double-repeats sites (n=48). **b,** Metaplot and heatmap showing enrichment of VDD, VAL, REM13, GDE1, CLSY3, CLSY4 and Pol IV ChIP-seq signal over single-repeat sites (n=37,456). **c,** Predicted aligned error (PAE) of VDD-VAL-GDE1-motif1. **d,** Interaction of GDE1 with VDD-VAL at three interfaces. Interacting residues and hydrogen bonds are highlighted in sticks and dashed lines, respectively.

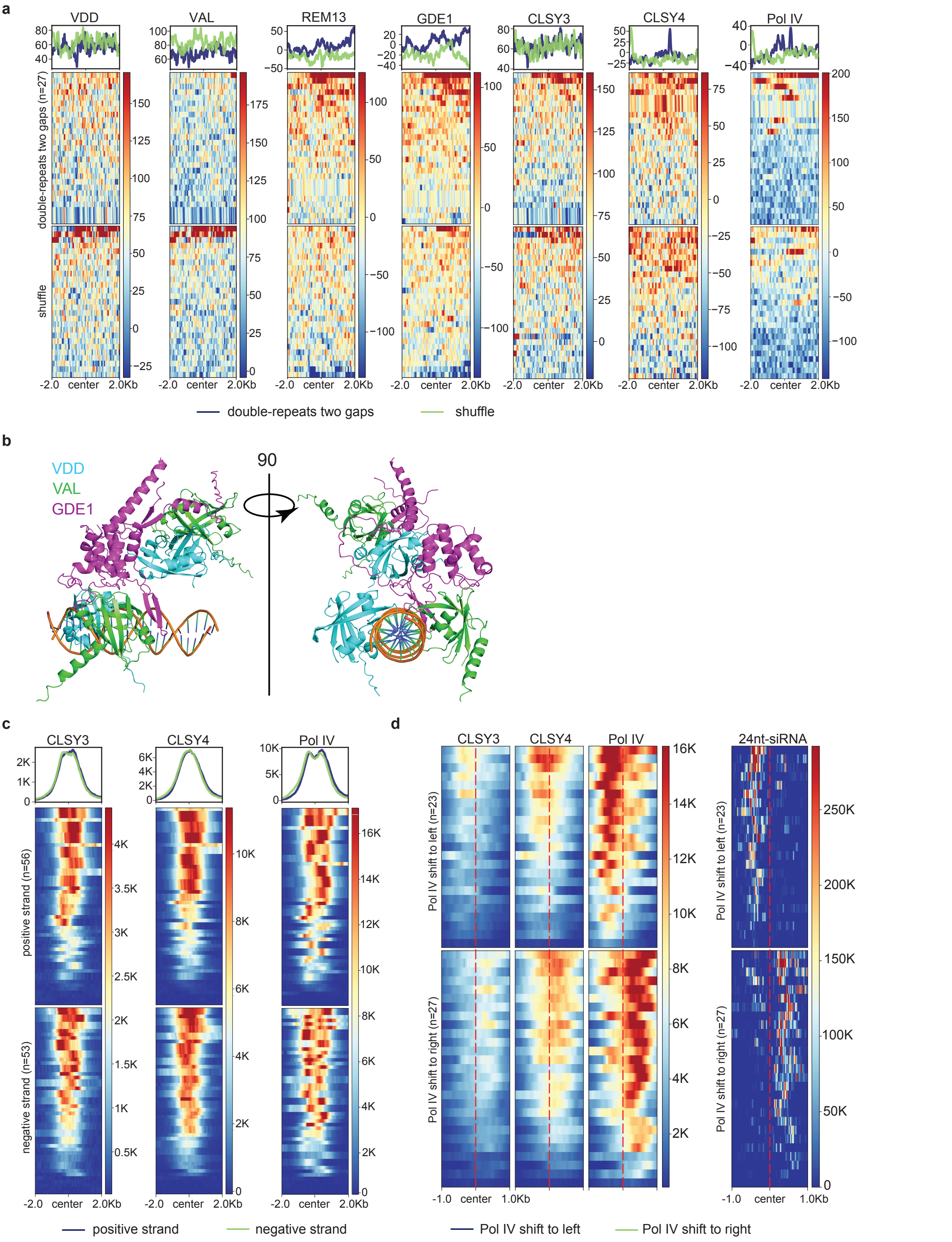


**Extended Data Figure 7 | Pol IV transcription complex localizes *clsy3 clsy4* motif1 for unidirectional transcriptions. a,** Metaplot and heatmap showing enrichment of VDD, VAL, REM13, GDE1, CLSY3, CLSY4 and Pol IV ChIP-seq signal over double-repeats with two gaps (n=27). **b,** Cartoon representation of VDD-VAL bound to dsDNA containing TTTTGCTTAT-GCTTTTGCTTAT. The DNA is shown as a ribbon representation. **c,** Metaplot and heatmap **showing** enrichment of CLSY3, CLSY4 and Pol IV ChIP-seq signal over all double-repeats sites at positive strand (n=56) or negative strand (n=53). **d,** Heatmap showing enrichment of CLSY3, CLSY4, Pol IV ChIP-seq signal and 24nt-siRNA signal over Pol IV left shift (n=23) or Pol IV right shift (n=27).

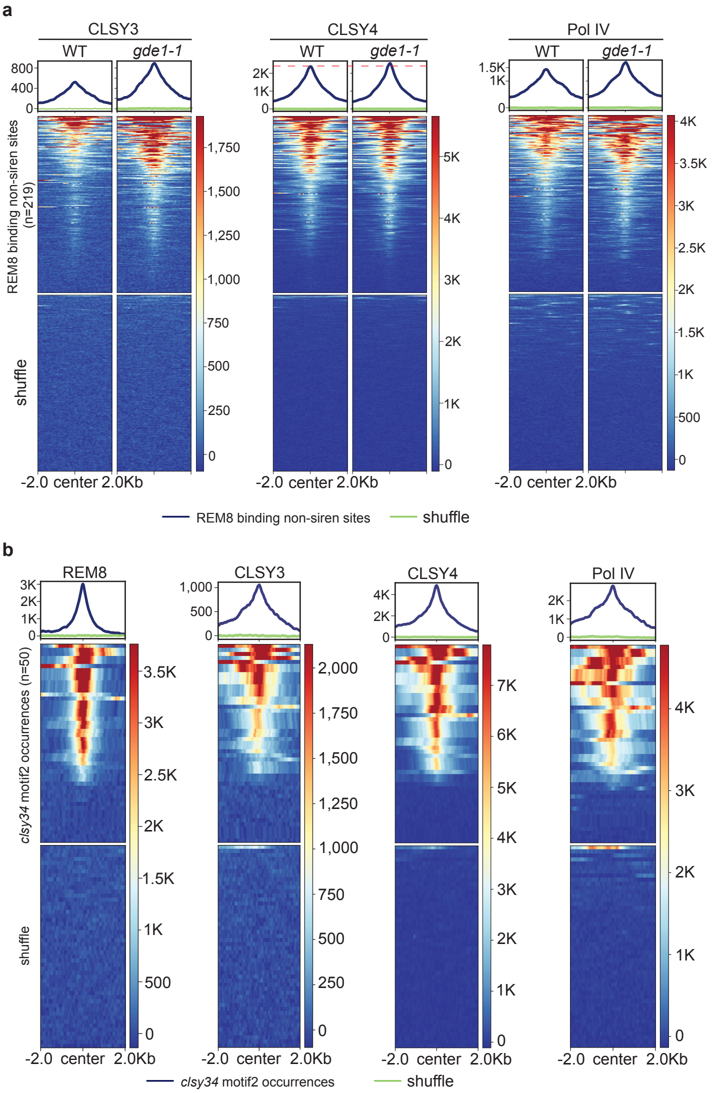


**Extended Data Figure 8 | Pol IV transcription complex recognizes *clsy3 clsy4* motif 2. a,** Metaplot and heatmap showing enrichment of CLSY3, CLSY4 and Pol IV ChIP-seq signal over REM8 binding non-siren sites and shuffle regions in WT or *gde1-1* backgrounds. **b,** Metaplot and heatmap showing enrichment of REM8, CLSY3, CLSY4, and Pol IV ChIP-seq signal over *clsy3 clsy4* motif2 sites (n=50).

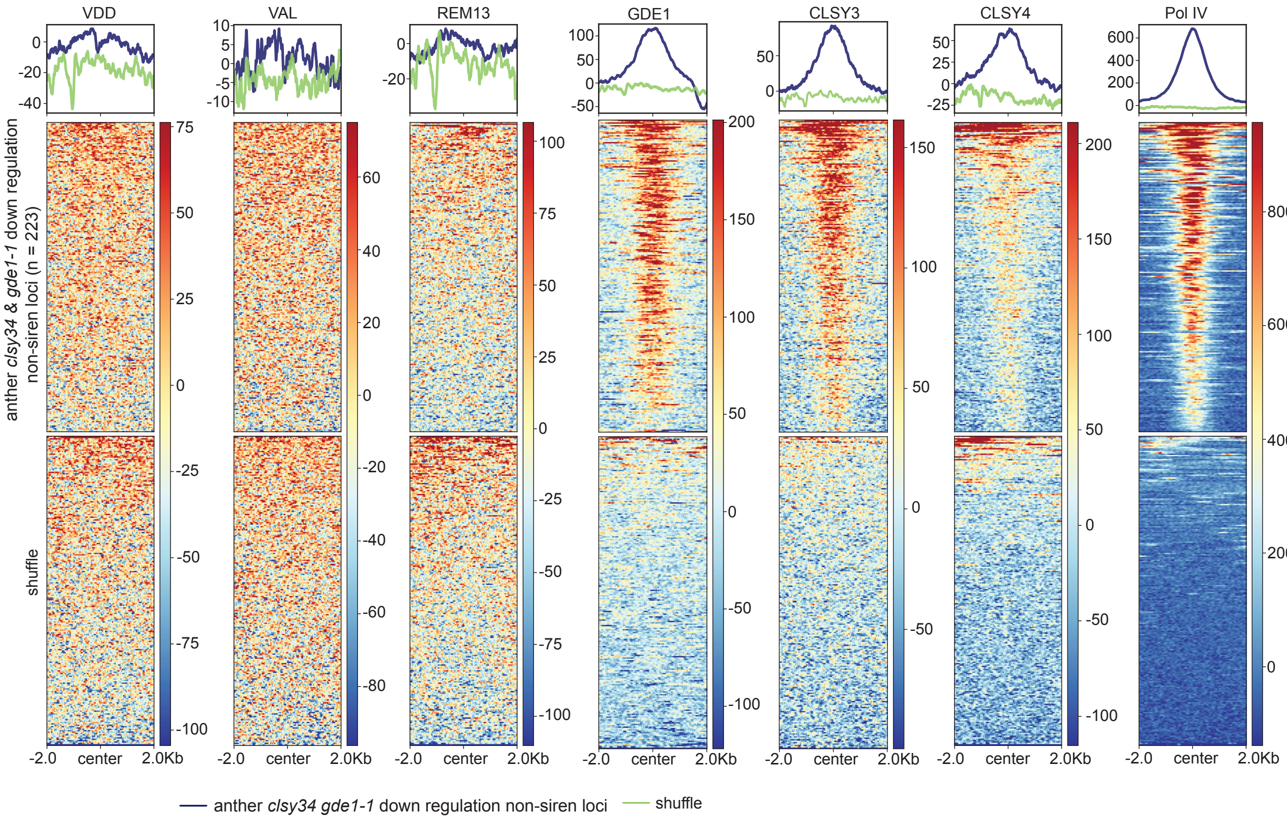


**Extended Data Figure 9 | GDE1, CLSY3/4 and Pol IV, but not REM transcription factors enriches at *clsy3 clsy4* and *gde1-1* down-regulated siRNA loci in anther.** Metaplot and heatmap showing enrichment of VDD, VAL, REM13, GDE1, CLSY3, CLSY4 and Pol IV ChIP-seq signal over anther *clsy3 clsy4* and *gde1-1* down regulation non-siren loci (n=223).
